# Root architecture plasticity in response to endoparasitic cyst nematodes is mediated by damage signaling

**DOI:** 10.1101/2022.06.07.495098

**Authors:** Nina Guarneri, Jaap-Jan Willig, Mark G. Sterken, Wenkun Zhou, M. Shamim Hasan, Florian M. W. Grundler, Viola Willemsen, Aska Goverse, Geert Smant, Jose L. Lozano-Torres

## Abstract

- Plant root architecture plasticity in response to biotic stresses has not been thoroughly investigated. Infection by the endoparasitic cyst nematodes induces root architectural changes that involve the formation of secondary roots at infection sites. However, the molecular mechanisms regulating secondary root formation in response to cyst nematode infection remain largely unknown.
- We first assessed whether secondary roots form in a nematode-density dependent manner by challenging wild type Arabidopsis plants with increasing numbers of cyst nematodes (*Heterodera schachtii*). Next, by using jasmonate-related reporter lines and knock-out mutants, we tested if tissue damage by nematodes triggers secondary root formation. Finally, we verified whether damage-induced secondary root formation depends on local auxin biosynthesis at nematode infection sites.
- Intracellular host invasion by *H. schachtii* triggers a transient local increase in jasmonates, which activates the expression of ERF109 in a COI1-dependent manner. Knock-out mutations in COI1 and ERF109 disrupt the nematode-density dependent increase of secondary roots observed in wildtype plants. Furthermore, ERF109 regulates secondary root formation upon *H. schachtii* infection via local auxin biosynthesis.
- Host invasion by *H. schachtii* triggers secondary root formation via the damage-induced jasmonate-dependent ERF109 pathway. This points at a novel mechanism underlying plant root plasticity in response to biotic stress.

## Introduction

Plants utilize root plasticity as a key strategy to survive in a changing soil environment. Remodeling of root systems allows plants to cope with nutrient deficiencies, drought, salinity, and other abiotic stresses (Koevoets *et al*., 2016). However, little is known about root architecture plasticity in response to soil-borne biotic stresses. Infections by cyst nematodes are known to induce elaborate root architectural changes in host plants. Secondary roots form locally at cyst nematode infection sites (Grymaszewska & Golinowski, 1991; Goverse *et al*., 2000; Lee *et al*., 2011).Furthermore, the ability to form secondary roots in response to nematode infection can result in better maintenance of shoot growth in some potato and soybean cultivars (Trudgill & Cotes, 1983; Miltner *et al*., 1991). Nevertheless, the molecular mechanisms regulating secondary root formation in response to belowground herbivory are not well understood.

Cyst nematodes are microscopic root endoparasites that cause large agricultural losses worldwide. These nematodes can persist in the soil in a dormant state for many years (Jones *et al*., 2013). Exudates from host roots trigger hatching of dormant second stage juveniles (J2s) and provide guidance for their migration to the root surface. Here, the J2s penetrate the root epidermis of the differentiation or mature root zone by piercing plant cell walls with their needle-like oral stylet and by secreting plant cell wall degrading enzymes (Bohlmann & Sobczak, 2014). Subsequently, juveniles migrate intracellularly within the cortex, leaving a trail of destruction (Wyss & Zunke, 1986; Grundler *et al*., 1994). Plant cell wall fragments released during nematode migration can act as damage-associated molecular patterns triggering defense signaling in the host (Shah *et al*., 2017). Nematode migration also activates biosynthesis and signaling of the defense hormone jasmonate (JA) (Kammerhofer *et al*., 2015). Upon successful arrival at the vascular cylinder, cyst nematodes utilize stylet-secreted effectors to manipulate plant developmental pathways to transform host cells into permanent feeding sites (Gheysen & Mitchum, 2011). Together with permanent feeding site development, multiple *de novo* formed secondary roots emerge in clusters at nematode infection sites (Grymaszewska & Golinowski, 1991; Goverse *et al*., 2000; Lee *et al*., 2011).

Nematode feeding sites are characterized by the local accumulation of the plant hormone auxin (Karczmarek *et al*., 2004; Grunewald *et al*., 2009). Auxin transport and auxin-insensitive Arabidopsis mutants infected by cyst nematodes show smaller females and smaller feeding sites, respectively (Goverse *et al*., 2000; Grunewald *et al*., 2009). Additionally, auxin is an important regulator of secondary root formation. Oscillations of auxin maxima at the root tip determine the formation of lateral roots in a regularly spaced pattern along the primary root (Fukaki & Tasaka, 2009). However, these oscillations are not required for the *de novo* formation of secondary roots. In fact, ectopic induction of local auxin biosynthesis in pericycle cells via an inducible promoter is sufficient to trigger *de novo* secondary root formation (Dubrovsky *et al*., 2008). Auxin accumulation in multiple neighboring pericycle cells can lead to the formation of secondary root clusters (Dubrovsky *et al*., 2008). The spatial co-occurrence of nematode feeding sites and secondary root clusters often corresponds to overlapping regions of auxin accumulation (Karczmarek *et al*., 2004; Absmanner *et al*., 2013). This suggests that secondary roots could be induced as the sole consequence of the auxin that accumulates during nematode feeding site development (Goverse *et al*., 2000). Alternatively, damage caused by nematode infection might also lead to local auxin accumulation and secondary root formation.

Tissue damage triggers auxin accumulation and *de novo* root formation via the JA-dependent ERF109 transcription factor in leaf explants (Liu *et al*., 2014; Chen *et al*., 2016; Hu & Xu, 2016; Zhang *et al*., 2019). Herein, JA accumulates at the site of wounding within a few hours of leaf detachment and triggers expression of the transcription factor ERF109 in a COI1-dependent manner. ERF109 binds to the promoter of the auxin biosynthesis gene ASA1, which induces root formation in a process referred to as *de novo* root organogenesis. Direct interaction of JAZ proteins inhibits ERF109 expression in a negative feedback loop to avoid wound hypersensitivity (Zhang *et al*., 2019). Sterile mechanical injury in primary roots of Arabidopsis can trigger auxin accumulation at the wounding site and subsequent secondary root formation (Sheng *et al*., 2017). However, it is unknown whether this occurs via the same damage signaling pathway as *de novo* root organogenesis from leaf explants. Furthermore, mechanical injury is an artificial condition and therefore it remains unclear whether the JA-dependent ERF109 pathway is involved in the regulation of secondary root formation also upon naturally occurring damage by herbivory or pathogen penetration.

Previously, we showed that components of the JA-dependent ERF109 pathway are induced by root-knot nematode (*Meloidogyne* spp.) infection (Zhou *et al*., 2019). Differently from cyst nematodes, root-knot nematodes penetrate roots at the elongation zone and migrate towards the root apical meristem by moving in between cells. Although this type of migration creates minimal tissue damage, root-knot nematode invasion of the root apical meristem induces expression of the ERF109 transcription factor. This eventually promotes tissue regeneration and reduces the inhibitory effect of nematode infection on primary root growth (Zhou *et al*., 2019). Thus, wound signaling can mediate primary root growth compensation in response to damage by stealthily migrating root-knot nematodes. However, further research is needed to understand whether JA-dependent wound signaling regulates root architectural changes to compensate for tissue destruction by the more damaging cyst nematodes in the differentiation and mature root zones.

In this study, we hypothesized that local tissue damage by cyst nematode host invasion causes secondary root formation at infection sites via the JA-dependent ERF109 pathway. By challenging Arabidopsis seedlings with increasing numbers of J2s of the beet cyst nematode *Heterodera schachtii*, we found that secondary root formation is induced at infection sites in a nematode-density dependent manner. With time course confocal microscopy of JA biosensors and ERF109 reporter lines in Arabidopsis, we provide evidence that secondary root formation is preceded by the transient and local JA-dependent expression of ERF109. Moreover, the nematode-density dependent increase in secondary roots is abolished in *coi1-2* and *erf109* knock-out mutants. By selectively applying the auxin biosynthesis chemical inhibitor L-kynurenine (L-kyn) to shoots and roots, we further found that the ERF109-mediated formation of secondary roots is dependent on local auxin biosynthesis. We therefore conclude that tissue damage by host invading cyst nematodes induces secondary root formation by altering local auxin biosynthesis via the JA-dependent ERF109 pathway. Altogether, our results show that damage signaling via the JA-dependent ERF109 pathway regulates root architectural plasticity in response to cyst nematode infection.

## Materials and methods

### Plant material and growth conditions

The Arabidopsis (*Arabidopsis thaliana*) lines Col-0, *pAOS::YFP_N_* (Poncini *et al*., 2017)*, DR5::GUS/Col-0 and DR5::GUS/erf109* (Cai *et al*., 2014), *p35S::JAS-VENUS/p35S::H2B-RFP* (Larrieu *et al*., 2015) and *pERF109::GFP/Col-0* (Zhou *et al*., 2019) were used. The *erf109* mutant was chosen because of the extensive characterization in previous research (Cai *et al*., 2014; Kong *et al*., 2018; Zhang *et al*., 2019; Ye *et al*., 2020). The weak allele *coi1-2* mutant (Xu *et al*., 2002) was used since it allows for propagation of homozygous plants and therefore, as opposed to other *coi1* mutant alleles, does not need pre-selection with MeJA, which could interfere with the ERF109 pathway. *pERF109::GFP/coi1-2* was obtained through crossing followed by selection of homozygous plants on selective ½ MS medium containing 15 µg ml^-1^ hygromycin B (Melford Laboratories Ltd.) and 20 µg ml^-1^ MeJA (Sigma-Aldrich). Arabidopsis plants were vertically grown in sterile conditions on modified Knop medium (Sijmons *et al*., 1991) in a growth chamber with a 16 h : 8 h, light : dark photoperiod at 21°C.

### Nematode sterilization

*Heterodera schachtii* (Woensdrecht population from IRS, the Netherlands) cysts were extracted from sand of *Brassica oleracea* infected plants as previously described (Baum *et al*., 2000) and incubated for seven days in a solution containing 1.5 mg ml^-1^ gentamycin sulphate, 0.05 mg ml^-1^ nystatin and 3mM ZnCl_2_. Hatched second stage juveniles (J2s) were purified by centrifugation on a 35% sucrose gradient and surface sterilized for 15 minutes in a solution containing 0.16 mM HgCl_2_, 0.49 mM NaN_3_, and 0.002% Triton X-100. After washing three times with sterile tap water, *H. schachtii* J2s were re-suspended in a sterile 0.7% Gelrite (Duchefa Biochemie, Haarlem, The Netherlands) solution. A similar concentration of Gelrite solution was used as mock treatment.

### Inoculation density-response curve

Individual Arabidopsis seeds were sown in 12×12cm square petri dishes. Nine-day old seedlings were inoculated with 0 (mock), 50, 100, 200, 350, or 500 *H. schachtii* J2s. Specifically, two 5 µl drops of solution (with J2s or mock) were pipetted at opposite sides of each seedling while keeping the petri dishes vertical. This allowed for a homogeneous smear of J2s along the whole length of the root. At 7 dpi, scans were made of whole seedlings using an Epson Perfection V800 photo scanner. The root architecture (total root length, primary root length, total secondary root length) was measured using the WinRHIZO package for Arabidopsis (WinRHIZO pro2015, Regent Instrument Inc., Quebec, Canada). For the *coi1-2* mutant, primary root length was measured manually because of the convoluted root system. The number of root tips was counted manually based on the scans. Furthermore, nematodes within the roots were stained with acid fuchsin and counted as previously described (Warmerdam *et al*., 2018). For comparisons between genotypes, the background effect of the mutation on the root architecture was corrected for by normalizing each measured component in infected seedlings to the average respective component in mock-inoculated roots. Additionally, the presence of clusters and the number of secondary root per cluster were scored using an Olympus SZX10 binocular.

### Histology and microscopy

Four-day-old Arabidopsis seedlings were inoculated with either 15 *H. schachtii* J2s or mock solution. The choice of using younger seedlings like previously done by Zhou *et al*. (2019) was made to reduce the possible amount of damage inflicted to the seedling during the preparation of the microscopic slides. Root architecture was inspected using an Olympus SZX10 binocular with a 1.5x objective and 2.5x magnification. Pictures were taken with an AxioCam MRc5 camera (Zeiss) and the ZEN 3.2 blue edition software (Zeiss). For confocal and brightfield microscopy, single-nematode infection sites were selected for observation. For histochemical staining of β-glucuronidase (GUS) activity, seedlings were incubated in a GUS staining solution (1 mg ml^-1^ X-GlcA in 100 mM phosphate buffer pH 7.2, 2 mM potassium ferricyanide, 2 mM potassium ferrocyanide, and 0.2 % Triton X-100) at 37°C (Zhou *et al*., 2019) for four hours. Stained seedlings were mounted in a chloral hydrate clearing solution (12 M chloral hydrate, 25% glycerol) and inspected with a Axio Imager.M2 light microscope (Zeiss) via a 20x objective. Differential interference contrast (DIC) images were taken with an AxioCam MRc5 camera (Zeiss) and the ZEN 3.2 blue edition software (Zeiss). GUS saturation was quantified as previously described (Beziat *et al*., 2017) using Fiji software (Schindelin *et al*., 2012). For confocal laser scanning microscopy, seedlings were mounted either in water or in 10 µg ml^-1^ propidium iodide (PI) and imaged using a Zeiss LSM 710 system via 10x and 40x objectives. The following wavelengths were used: 600–640 nm for PI, 500-540 nm for GFP, 520-560 nm for YFP, and 590-680 nm for RFP. For *pAOS_N_::YFP* and *JAS9-VENUS* reporters, the fluorescent signal was imaged at the focal plane displaying the xylem vessels, where the nematode head is found. For the *pERF109::GFP* reporter, Z-stacks of six 13μm-slices were made of the entire root depth. Images were taken using the ZEN 2009 software (Zeiss) and processed using Fiji software. To make the fluorescence more visible, the brightness was enhanced for all the representative pictures in the same way using Adobe Photoshop 2021. Fluorescence intensity was quantified using the Fiji software. Specifically, the region of interest was selected using a set threshold and then the integrated density was measured. Z-Stacks were projected using the maximum intensity method.

### Auxin biosynthesis inhibition

For split-plate assays, we used the method as described in Matosevich *et al*. (2020). For the L-kyn split plate assay the following four treatment combinations were prepared: MM (modified Knop medium and 0.02% DMSO), KK (modified Knop medium, 10 µM L-kynurenine (Sigma-Aldrich) and 0.02% DMSO), MK (L-kyn only in the root), and KM (L-kyn only in the shoot). For the Yuc split plate assay the following four treatment combinations were prepared: MM (modified Knop medium and 0.2% DMSO), YY (modified Knop medium, 100 µM Yucasin (Sigma-Aldrich) and 0.2% DMSO), MK (Yuc only in the root), and YM (Yuc only in the shoot). Four-day-old Arabidopsis seedlings were inoculated with 15 *H. schachtii* J2s. Sixteen hours post inoculation, when J2s are still migrating through the root, seedlings were transferred to the treatment plates, so that the shoot and the hypocotyl were in contact with the medium in the upper half of the plate and the nematode-infected root was on the medium in the lower half of the plate. For microscopy, seedlings were collected at 3 dpi and GUS staining was performed (as described above). For root architecture inspection, scans were made of whole seedlings at 7 dpi using an Epson Perfection V800 photo scanner. The total number of secondary roots per plant was counted based on the scans. Additionally, the presence of clusters and the number of secondary roots per cluster were scored using an Olympus SZX10 binocular.

### Statistical analyses

Statistical analyses were performed using the R software version 3.6.3 (Windows, x64). The R packages used were tidyverse (https://CRAN.R-project.org/package=tidyverse), ARTool (https://CRAN.R-project.org/package=ARTool) and multcompView (https://CRAN.R-project.org/package=multcompView). Correlation between variables was calculated using Spearman Rank-Order Correlation coefficient. For binary data, significance of the differences between proportions was calculated by a Pairwise Z-test. For normally distributed data, significance of the differences among means was calculated by Student’s T-test for pairwise comparisons and by ANOVA followed by Tukey’s HSD test for multiple comparisons. A non-parametric pairwise Wilcoxon test followed by false discovery rate correction for multiple comparisons was used for non-normally distributed data with one grouping factor. For non-parametric two-factorial ANOVA, an Aligned Rank Transform followed by Tukey’s HSD test for multiple comparisons was applied. The confidence interval of the inoculum density-response curves was calculated by loess regression (as per default in geom_smooth) in R.

## Results

### *H. schachtii* infection induces local formation of secondary roots in a nematode density-dependent manner

To test if tissue damage by invading nematodes in roots triggers the formation of secondary roots, we analyzed root branching upon penetration by increasing numbers of nematodes. We inoculated seedlings with 0, 50, 100, 200, 350, or 500 J2s of *H. schachtii* and counted both the number of nematodes that penetrated the roots and the total number of secondary roots at 7 dpi (Fig. 1). Here, the total number of secondary roots in infected seedlings was normalized to the average respective number in uninfected roots. We found that the number of nematodes that penetrated the roots increased by inoculum density for up to 350 J2s per plant, whereafter it remained the same (Fig. 1a). Furthermore, we observed that the number of penetrated nematodes correlated positively with the total number of secondary roots per plant (Fig. 1b). Next, we investigated if the clustering of secondary roots around nematode infections sites also correlates with the inoculum density (Fig. 1c,d). For this, we challenged Arabidopsis with four inoculum densities (0, 50, 100, and 350) to establish an incremental increase in the number of nematode infection sites per plant. Infection sites were identified by the local discoloration of root tissue due to cell necrosis along the migratory tract of the nematode (Grundler *et al*., 1994). Roots were counted as clusters when more than one secondary root emerged in proximity of an infection site. Also, we counted the number of secondary roots per cluster. We found that uninfected seedlings showed a typical pattern of lateral roots regularly distributed along the primary root (Fig. 1c). However, in infected seedlings clusters of secondary roots emerged close to nematode infection sites in an inoculum density-dependent manner (Fig. 1c,d). Interestingly, also the number of secondary roots per cluster significantly increased at inoculum density 350 compared to 50 (Fig. 1c,e). Moreover, higher inoculation densities caused more extensive discoloration at the infection sites indicating higher levels of tissue damage. Altogether, these observations showed that infection by *H. schachtii* triggers local density-dependent formation of secondary roots.

**Fig. 1.**
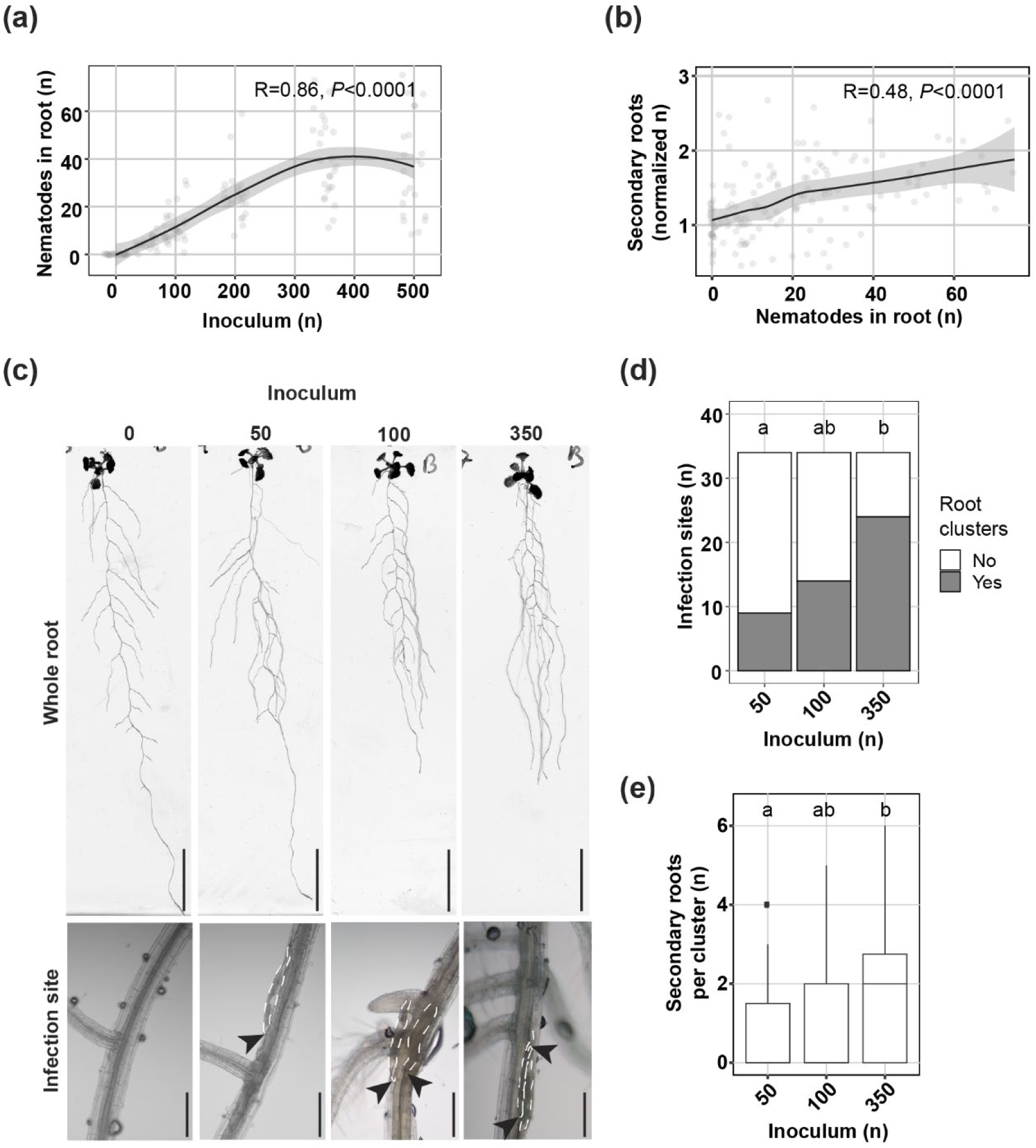
Secondary roots form locally at *Heterodera schachtii* infection sites in a nematode density-dependent manner. Nine-day-old Arabidopsis Col-0 seedlings were inoculated with increasing numbers of *H. schachtii* J2s, ranging from 0 (mock) to 500 J2s per seedling. At 7 days post inoculation (dpi), scans were made of the root systems and the total number of secondary roots per plant was counted. Fuchsin staining was performed to count the number of J2s that had penetrated the roots. Additionally, the presence of clusters and the number of secondary roots per cluster was scored. (a) Number of nematodes that successfully penetrated the roots per inoculum. (b) Number of secondary roots formed per number of nematodes inside the roots. Total number of secondary roots in infected seedlings was normalized to the average respective component in uninfected roots and correlated with the number of nematodes inside the roots. Data from three independent biological repeats of the experiment was combined. Correlation (R) between two variables was calculated using Spearman’s Rank-Order Correlation coefficient (n=30, *P*<0.0001). Grey area indicates the 95% confidence interval. (c) Representative pictures of whole roots and infection sites in Col-0 seedlings inoculated with 0 (mock), 50, 100, and 350 J2s. Scale bars in whole root and infection site pictures are 2 cm and 200 µm, respectively. Black arrowheads indicate the nematodes head; white dotted lines outline the nematodes body. (d) The proportions of secondary root clusters close to infection sites in Arabidopsis seedlings inoculated with 50, 100, and 350 J2s. Statistical significance was calculated by a Pairwise Z-test (n=34, p<0.05). (e) Number of secondary roots within each root cluster in Arabidopsis seedlings inoculated with 50, 100, and 350 J2s. Statistical significance was calculated by pairwise Wilcoxon test followed by false discovery rate correction for multiple comparisons (n=34, p<0.001). Different letters indicate statistically different groups.

### *H. schachtii* host invasion induces JA biosynthesis and signaling

Artificially induced tissue damage can trigger the formation of roots via JA-dependent signaling pathways. For instance, wounding induces JA-dependent *de novo* root organogenesis in leaf explants (Zhang *et al*., 2019). Infective juveniles of *H. schachtii* invade the host by destructive thrusts of the oral stylet and release of plant cell wall degrading enzymes causing extensive cell damage during host invasion (Grundler *et al*., 1994; Tytgat *et al*., 2002; Vanholme *et al*., 2007). We hypothesized that secondary root formation in proximity of *H. schachtii* infection sites might be regulated by JA, in response to tissue damage associated with nematode host invasion. To test our hypothesis, we investigated whether JA biosynthesis and signaling were activated during *H. schachtii* infection using the JA biosynthesis reporter line *pAOS::YFP_N_* (Poncini *et al*., 2017) and the JA signaling biosensor *JAS9-VENUS* (*p35S::JAS9-VENUS/p35S::H2B-RFP*) (Larrieu *et al*., 2015) (Fig. 2). We chose three timepoints that reflect the early parasitic stages of intracellular host invasion (12 hours post inoculation, hpi), permanent feeding site initiation (24 hpi), and permanent feeding site expansion (168 hpi) (Tytgat *et al*., 2002; Hewezi *et al*., 2014; Kammerhofer *et al*., 2015; Marhavy *et al*., 2019). Importantly, to avoid interference of signals due to the presence of multiple nematodes at one infection site, we selected single-nematode infection sites for our observations. We found that infection with *H. schachtii* significantly induces transient expression of *pAOS::YFP_N_*, with the highest level of expression at 12 hpi (Fig. 2a-d). Likewise, *JAS9-VENUS* showed a strong JA signaling activity (i.e., low VENUS/RFP ratio) in infected roots at 12 hpi, which decreased over time to the level of uninfected root tissue at 168 hpi (Fig. 2e-h). These observations demonstrated that both JA biosynthesis and JA signaling are strongly induced during and shortly after *H. schachtii* host invasion close to the nematode infection site. We therefore concluded that tissue damage caused by *H. schachtii* during intracellular host invasion triggers local JA biosynthesis and signaling in Arabidopsis.

**Fig. 2.**
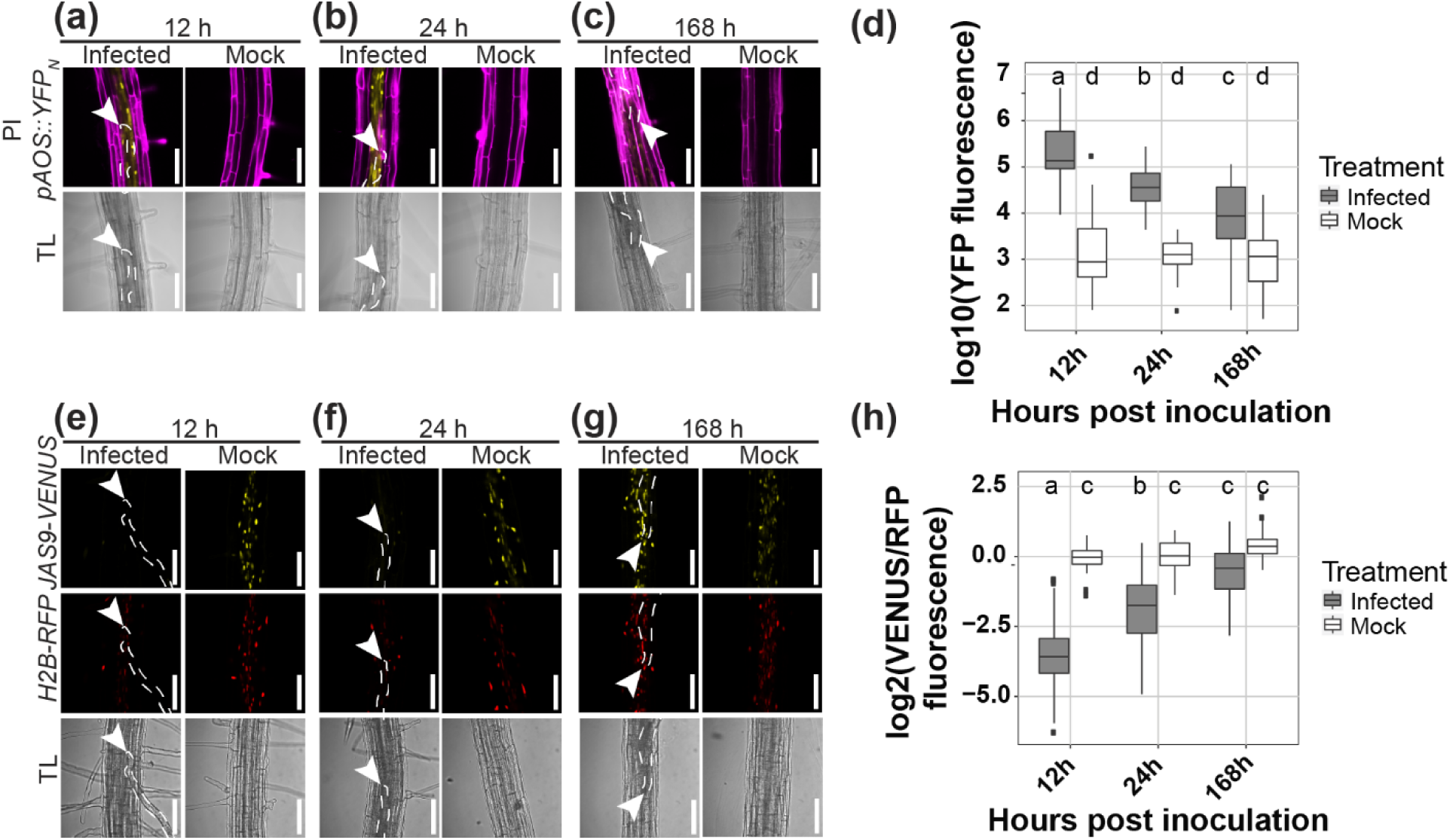
Transient induction of JA biosynthesis and signaling at *Heterodera schachtii* infection sites. Four-day-old Arabidopsis seedlings were either inoculated with 15 *H. schachtii* J2s or mock inoculated. At 12, 24, and 168 hpi seedlings were mounted in 10 μg ml^-1^ propidium iodide (PI) and then imaged using a fluorescent confocal microscope. Single-nematode infection sites were selected for observation. (a-c) Representative pictures of infected and non-infected roots expressing the JA biosynthesis marker *pAOS::YFPN*. To make the fluorescence more visible, the brightness was enhanced for all the representative pictures in the same way. (d) Quantification of YFP intensity in the *pAOS::YFPN* line. Values represent the log10 of the YFP integrated density. (e-g) Representative pictures of infected and non-infected roots expressing the JA biosensor *p35S::JAS9-VENUS/p35S::H2B-RFP*. To make the fluorescence more visible, the brightness was enhanced for all the representative pictures in the same way. (h) Quantification of the JA signaling repressor motif *JAS9*. Values represent the log2 of the fluorescence ratio between *JAS9-VENUS* and *H2B-RFP* raw integrated densities. Data from three independent biological repeats of the experiment was combined. Significance of differences between fluorescent intensities in nematode-infected and non-infected seedlings over the different timepoints was calculated by ANOVA followed by Tukey’s HSD test for multiple comparisons (n=30, *P*<0.0001). Different letters indicate statistically different groups. White arrowheads indicate the nematode head; white dotted lines outline the nematode body. TL=transmission light. Scale bar is 100 µm.

### COI1-mediated JA signaling regulates *ERF109* expression upon *H. schachtii* infection

Root tip resection or wounding in leaf explants induce *ERF109* expression in a COI1-dependent manner (Zhang *et al*., 2019; Zhou *et al*., 2019). To determine if *H. schachtii-*induced JA signaling also triggers *ERF109* expression, we monitored *pERF109::GFP* expression within single-nematode infection sites in the *coi1-2* mutant and wildtype Arabidopsis Col-0 plants during the early stages of infection by *H. schachtii* (Fig. 3). Similar to what we observed for JA biosynthesis and signaling, *ERF109* expression was induced at early timepoints (12 and 24 hpi) of *H. schachtii* infection around the migratory track of the nematodes in wildtype Col-0 (Fig. 3a-c). Moreover, in the *coi1-2* mutant, *pERF109::GFP* fluorescence was significantly reduced compared to wildtype Arabidopsis (Fig. 3d). We nonetheless observed a slight increase in GFP fluorescence in *coi1-2* mutant over time, which might be caused by tissue autofluorescence from the cell walls of the maturing permanent feeding sites (Hoth *et al*., 2005). Since *pERF109::GFP* has a nuclear-cytoplasmic localization (Zhou *et al*., 2019), autofluorescence from cell walls in syncytia cannot be easily distinguished from the cytoplasmic part of the *pERF109::GFP* signal. However, after quantifying only nuclear-localized *pERF109::GFP*, the initially observed increase in fluorescence in the nematode-infected *coi1-2* mutant over time was not detected anymore, pointing at autofluorescence as the most plausible cause for the increasing GFP signal in maturing permanent feeding sites (Fig. S1). We therefore concluded that *H. schachtii* induces *ERF109* expression during host invasion in a JA-dependent manner.

**Fig. 3.**
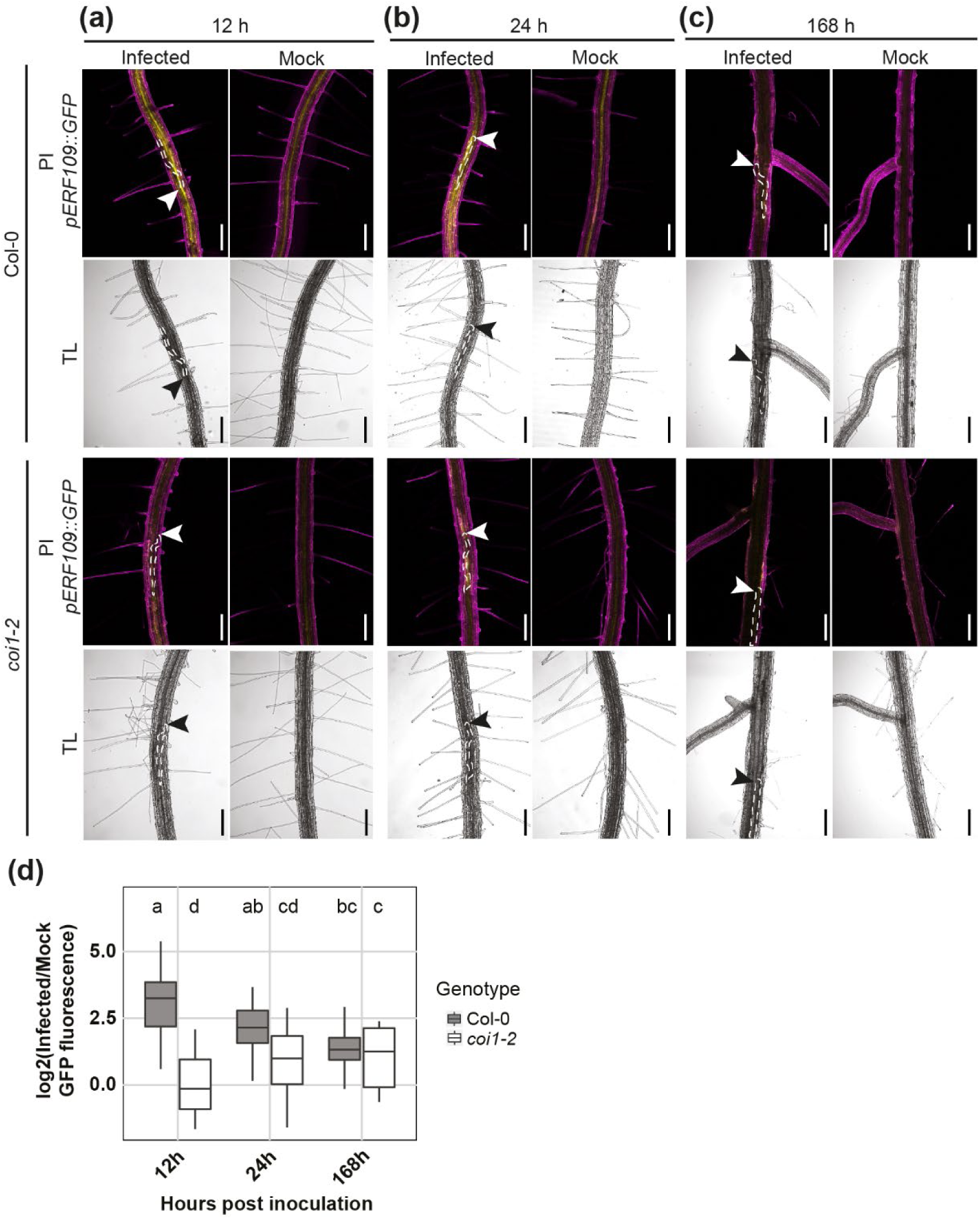
*ERF109* expression upon *Heterodera schachtii* host invasion is dependent on COI1-mediated JA signaling. Four-day-old Arabidopsis seedlings were either inoculated with 15 *H. schachtii* J2s or mock inoculated. At 12, 24, and 168 hpi seedlings were mounted in 10 μg ml^-1^ propidium iodide (PI) and then imaged using a fluorescent confocal microscope. Single-nematode infection sites were selected for observation. (a-c) Representative pictures of infected and mock-inoculated seedlings expressing the *pERF109::GFP* construct in either wildtype Col-0 or mutant *coi1-2* background at 12 hpi (a), 24 hpi (b), and 168 hpi (c). To make the fluorescence more visible, the brightness was enhanced for all the representative pictures in the same way. (d) Quantification of *pERF109::GFP* fluorescent intensity induced by infection of Col-0 and *coi1-2* roots. Values represent log2 of the fluorescence ratio between the GFP integrated density of infected and non-infected roots. Data from two independent biological repeats of the experiment was combined. Significance of differences between fluorescent intensities in Col-0 and *coi1-2* roots over the different timepoints was calculated by ANOVA followed by Tukey’s HSD test for multiple comparisons (n=20, *P*<0.05). Different letters indicate statistically different groups. White and black arrowheads indicate the nematode head; white dotted lines outline the nematode body. TL=transmission light. Scale bar is 200 µm.

### COI1 and ERF109 regulate secondary root formation upon *H. schachtii* infection

Next, we asked if the activation of JA-dependent expression of *ERF109* is required for the formation of secondary roots during *H. schachtii* infections. If this holds true, the nematode density-dependent increase in secondary roots observed for wildtype Col-0 should be altered in both *coi1-2* and *erf109* mutants. To test this, we performed the same density-response experiment as shown in figure 1a and b. At 7 dpi, the number of nematodes that had successfully penetrated the roots did not differ significantly between wildtype Col-0 and the *erf109* mutant (Fig. 4). In contrast, the number of nematodes was significantly higher in roots of the *coi1-2* mutant compared to wildtype Arabidopsis plants, indicating a role of COI1 in plant susceptibility to penetration by *H. schachtii* (Fig. 4b). However, it must be noted that the uninfected *coi1-2* mutant had a much larger root system compared to wildtype Arabidopsis Col-0 (Fig. S4), which also may influence the number of nematode penetrations. Nevertheless, while nematode infections in wildtype Arabidopsis induced the formation of secondary roots, no such increase was observed for *erf109* and *coi1-2* mutants (Fig. 4c). In conclusion, both COI1 and ERF109 regulate the density-dependent induction of secondary root formation by *H. schachtii*. This induction of secondary root formation is independent from plant susceptibility to nematode penetration.

**Fig. 4.**
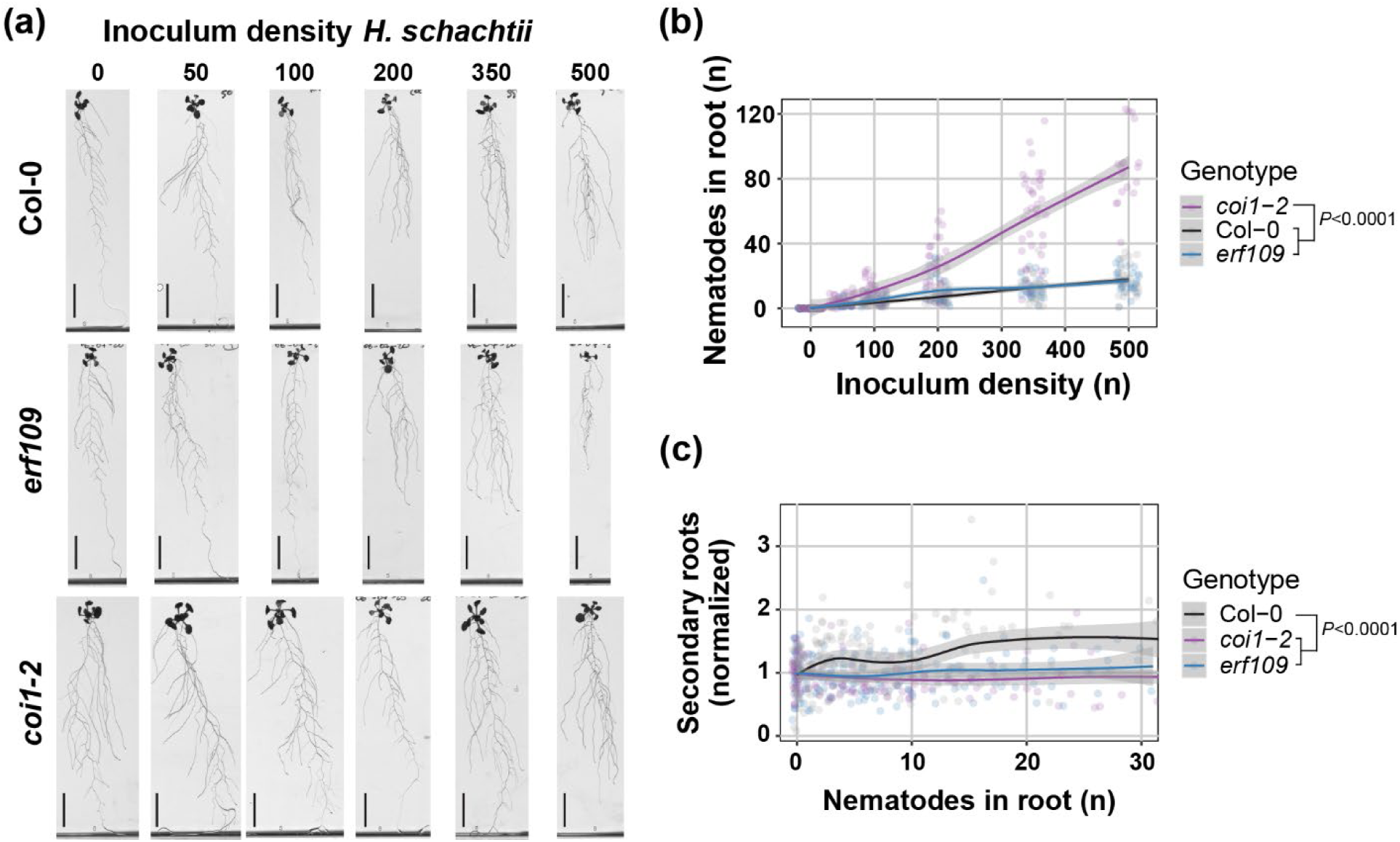
ERF109 and COI1 regulate the nematode density-dependent secondary root formation that is triggered by *Heterodera schachtii* infections. Nine-day-old wildtype Col-0, *erf109,* and *coi1-2* seedlings were inoculated with increasing numbers of *H. schachtii* J2s, ranging from 0 (mock) to 500 J2s per seedling. At 7 dpi, scans were made of the root systems and the number of secondary roots per plant was counted. Fuchsin staining was performed to count the number of J2s that had penetrated the roots. (a) Representative pictures of wildtype Col-0, *erf109* and *coi1-2* infected seedlings at 7 dpi. (b) Number of nematodes that successfully penetrated the roots per inoculum. (c) Secondary roots formed per number of nematodes inside the roots. Total number of secondary roots in infected seedlings was normalized to the average respective component in mock-treated roots. Data of three independent biological repeats of the experiment was combined. Significance of differences between genotypes was calculated by ANOVA followed by Tukey’s HSD test for multiple comparisons (n=30, *P*<0.0001). Grey area indicates the 95% confidence interval. Scale bar is 2 cm.

### ERF109-mediated induction of secondary root formation compensates for primary root growth inhibition by *H. schachtii*

The induction of secondary root formation by cyst nematodes might compensate for a possible inhibition of root growth by nematode invasion. To test this hypothesis, we investigated whether the total length of the entire root system, the primary root length, and the total length of the secondary roots were altered in the infected *erf109* mutant compared to wildtype Col-0 (Fig. 5). To eliminate the background effect of the mutation on the root architecture, we normalized each measured component in infected seedlings to the average respective component in uninfected roots. We found that the total length of the root system of wildtype Col-0 at increasing numbers of nematodes remains similar to that of uninfected plants (i.e., close to 1 in Fig. 5a). In contrast, the total length of the entire root system in the *erf109* mutant decreased by nematode density as compared to uninfected plants. As the total length of the root system is the sum of the lengths of the primary roots and the secondary roots, we also analyzed these components separately. The primary root length of both wildtype Col-0 and the *erf109* mutant declined by nematode density (Fig. 5b). This decline was slightly but significantly exacerbated by the *erf109* mutation. However, we found a more striking difference in the total length of the secondary roots between wildtype Col-0 and the *erf109* mutant (Fig. 5c). In wildtype Col-0, we observed a significant increase in the total length of the secondary roots by nematode density, sufficient to compensate for the loss in primary root length. However, we observed no significant increase in the total length of the secondary roots by nematode density in the *erf109* mutant, which explains why the total length of the root system by nematode density remained stable for wildtype Col-0, but not for the *erf109* mutant. Based on our data, we conclude that ERF109-mediated formation of secondary roots compensates for primary root growth inhibition by *H. schachtii*.

**Fig. 5.**
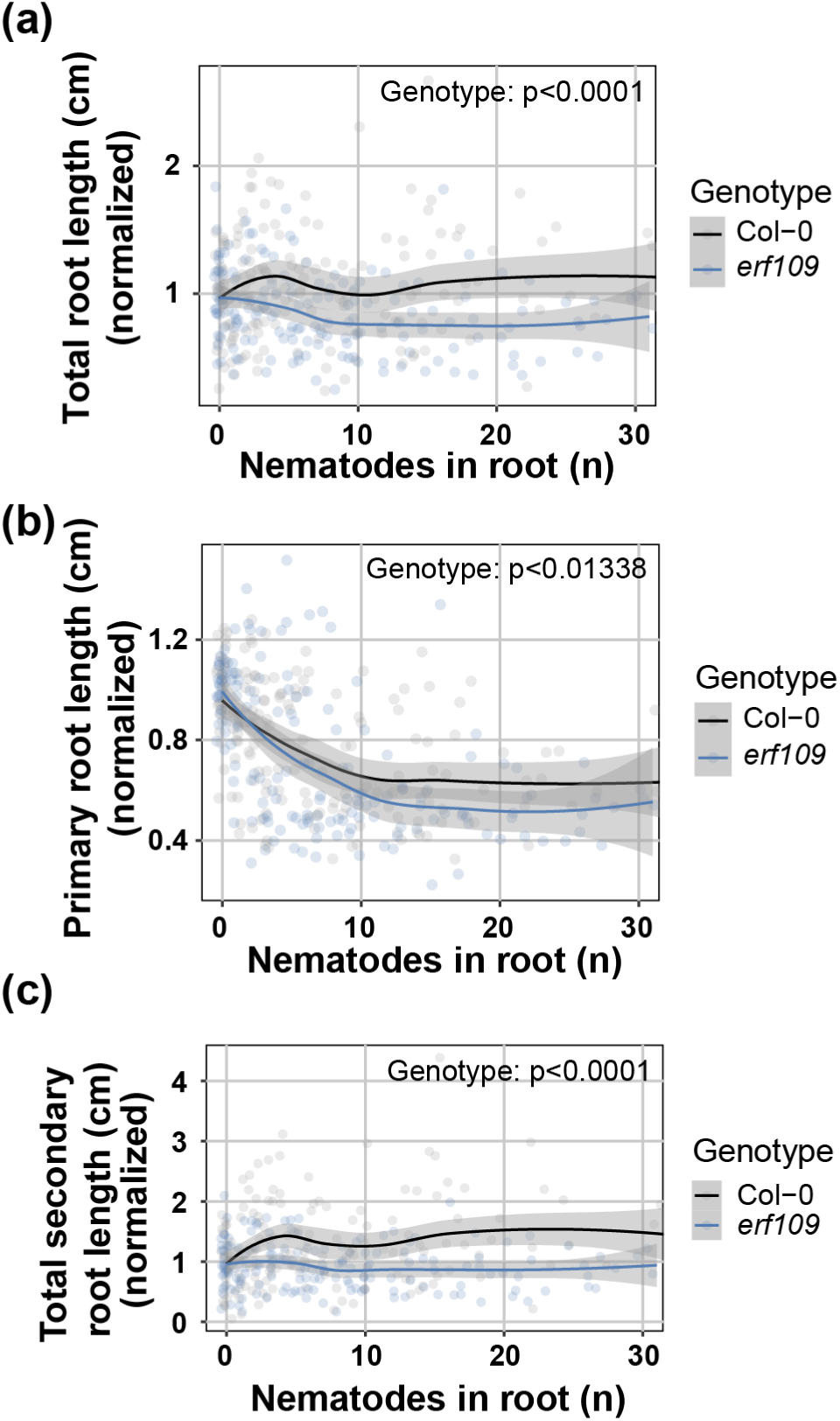
ERF109-mediated secondary root formation allows for maintenance of total root length despite primary root growth inhibition by *Heterodera schachtii*. Nine-day-old Col-0 and *erf109* Arabidopsis seedlings were inoculated with increasing *H. schachtii* densities ranging from 0 (mock) to 500 J2s per seedling. At 7 dpi, scans were made of the root systems and the root length was measured using WinRHIZO. Total, primary, and secondary root length was normalized to the average respective component in mock-treated roots. Fuchsin staining was performed for counting the number of J2s that penetrated the roots. (a) Total root length per number of nematodes in the roots. (b) Primary root length per number of nematodes in the roots. (c) Total secondary root length per number of nematodes in the roots. Data from three independent biological repeats of the experiment was combined. Significance of differences between genotypes was calculated by ANOVA (n=30). Grey area indicates the 95% confidence interval.

### ERF109 regulates local auxin biosynthesis at the nematode infection site

ERF109 mediates JA-induced secondary root formation by directly binding to the promoter of auxin biosynthesis genes *ASA1* and *YUC2* (Cai *et al*., 2014). We hypothesized that ERF109 regulates secondary root formation by inducing local auxin biosynthesis at the nematode infection site. Thus, we used a split plate assay containing growth media with and without L-kyn to chemically inhibit auxin biosynthesis in the shoots and/or the roots of infected wildtype and *erf109* plants (Fig. 6). The local accumulation of auxin was monitored using the *DR5::GUS* reporter (Fig. 6a). When seedlings were grown on regular medium or when auxin biosynthesis was inhibited by L-kyn only in the shoots, *DR5::GUS* was expressed at nematode infection sites in wildtype Col-0 seedlings. However, when auxin biosynthesis was inhibited in both shoots and roots or only in the roots by treatment with L-kyn, no *DR5::GUS* expression was observed (Fig. 6b,c). This suggested that auxin accumulation at nematode infection sites was dependent on local auxin biosynthesis in the roots. Importantly, we observed that the auxin accumulation at nematode infection sites via root-localized auxin biosynthesis was disrupted in the *erf109* mutant. Indeed, *DR5::GUS* expression was significantly lower at the nematode infection sites in *erf109* seedlings compared to wildtype Col-0 when auxin biosynthesis was permitted in the root (Fig. 6b,c). To determine if the differences in *DR5::GUS* between the two Arabidopsis genotypes were only local at the nematode infection site or systemic throughout the root system, we also looked at *DR5::GUS* expression in root tips (Fig. 6d, S2). In contrast to nematode infection sites, when auxin biosynthesis was inhibited only in the shoots, we observed no difference between *erf109* and wildtype Col-0 in *DR5::GUS* expression in the root tip (Fig. 6d, S2). Since L-kyn has been shown to also inhibit ethylene-induced auxin biosynthesis (He *et al*., 2011) we also performed the experiment using the auxin biosynthesis inhibitor Yucasin (Yuc). Due to the higher concentration of DMSO used to dissolve Yuc, an overall lower frequency of *DR5:GUS* staining was observed. Nevertheless, the Yuc split plate assay showed the same trend as the L-kyn experiment (Fig. S4). From these results, we concluded that ERF109 regulates local auxin biosynthesis at infection sites of *H. schachtii*.

**Fig. 6.**
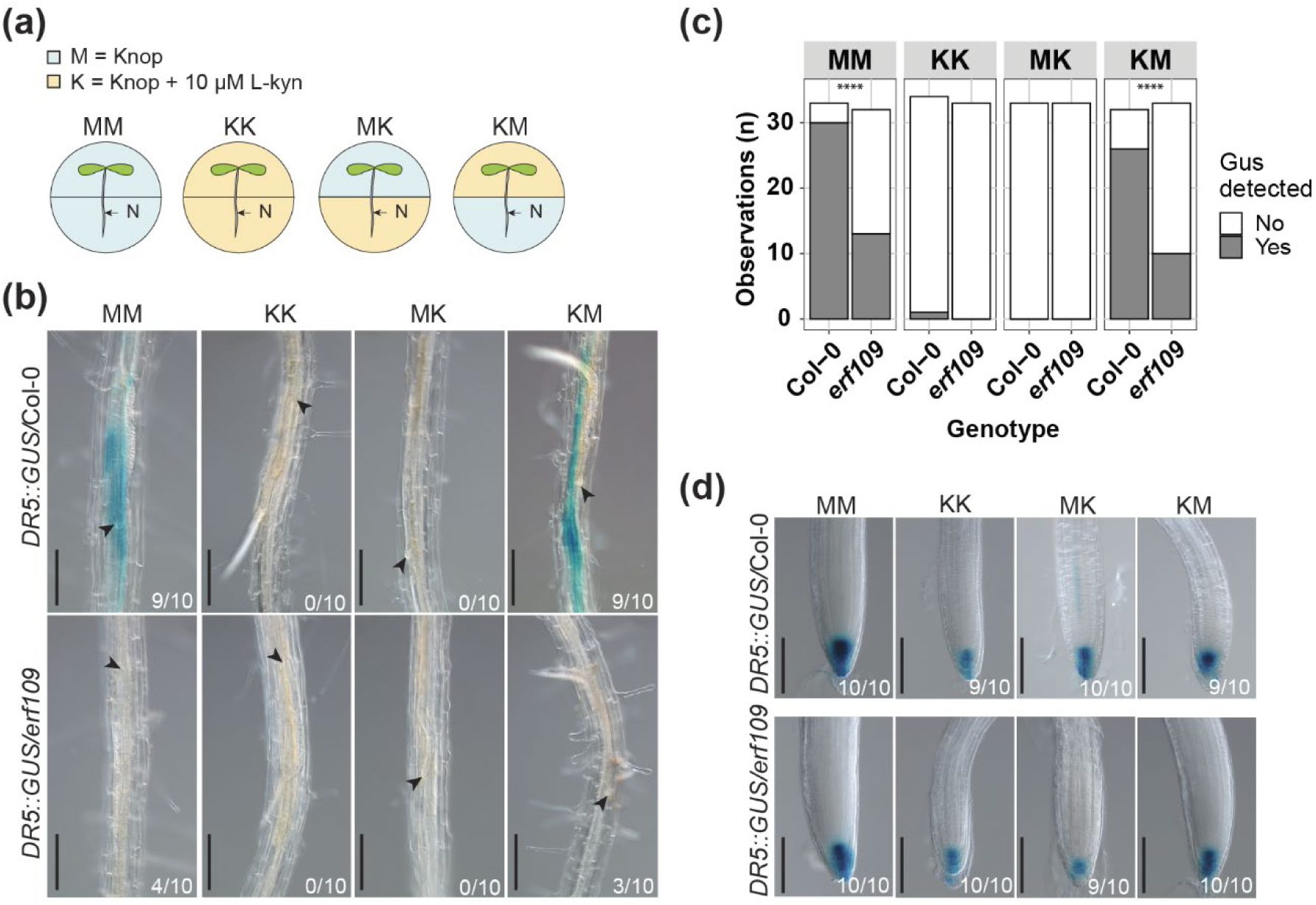
ERF109 regulates local auxin biosynthesis at *Heterodera schachtii* infection sites. Four-day-old Arabidopsis Col-0 and *erf109* seedlings expressing the auxin *DR5::GUS* reporter were infected with 15 *H. schachtii* J2s. At 16 hours post inoculation, seedlings were transferred to treatment plates. Four treatment combinations were prepared: MM (modified Knop medium and 0.02% DMSO), KK (modified Knop medium, 10µM L-kyn and 0.02% DMSO), MK (L-kyn only in the root), KM (L-kyn only in the shoot). At 3 dpi GUS staining assay was performed for 4 hours and seedlings were imaged. Single-nematode infection sites were selected for observation. (a) Experimental design with Arabidopsis seedlings transferred to split plates with modified Knop medium either with or without L-kyn. N = nematode. (b) *DR5::GUS* expression at nematode infection sites in wildtype Col-0 and *erf109* roots in the four different treatment combinations with or without L-kyn applied to shoots and/or roots. (c) Number of observations with (Yes) or without (No) GUS staining at the nematode infection sites in roots of wildtype Col-0 and *erf109* plants. Statistical significance was calculated by a Pairwise Z-test (n=33, ****, *P*<0.0001). (d) *DR5::GUS* expression in the root tips of Col-0 and *erf109* roots. Black arrowheads indicate the nematode head. Frequencies at the bottom right corner indicate how many times GUS staining was observed in one of the three independent biological repeats of the experiment. Scale bar is 200 µm.

### ERF109-induced secondary root formation upon *H. schachtii* infection is dependent on local auxin biosynthesis

We found that ERF109 regulates local auxin biosynthesis at *H. schachtii* infection sites. This raised the question if the ERF109-mediated secondary root formation upon *H. schachtii* infection is dependent on this local biosynthesis of auxin. To test this, we inoculated four-day-old wildtype Col-0 and *erf109* seedlings with either 15 *H. schachtii* J2s or a mock solution. At 16 hpi, seedlings were transferred to the four previously described split-plates containing medium with and without 10 µM L-kyn (Fig. 6a). At 7 dpi, the total number of secondary roots was scored. As expected, the different treatment combinations with and without L-kyn in the shoots and/or roots led to a different number of lateral roots in the uninfected roots (Fig. S3). Therefore, to calculate the number of additional secondary roots induced by nematode infection, the number of secondary roots in infected roots was normalized to the average respective component in uninfected roots. Additionally, we scored how often a cluster of roots occurs in proximity of an infection site and the number of secondary roots per cluster (Fig. 7). When auxin biosynthesis was inhibited in both shoots and roots or only in the roots no additional secondary roots formed in infected Col-0 wildtype seedlings (Fig. 7a,b). Consistently, no clusters of secondary roots were found at nematode infection sites (Fig. 7c-e). However, inhibition of auxin biosynthesis in the shoots alone led to a significant reduction in the total number of secondary roots in infected seedlings (Fig. 7a,b) as well as in the number of clusters and in the number of secondary roots per cluster compared to when auxin biosynthesis was permitted in both shoots and roots (Fig. 7c-e; treatment MM versus KM). Thus, secondary root formation upon *H. schachtii* infection is dependent on local auxin biosynthesis, although polar auxin transport from the shoots might still play a role. Furthermore, the mutation in *erf109* strongly affected secondary root formation when auxin biosynthesis was permitted in the roots. Indeed, a significant decrease in the number of additional secondary roots, in the number of clusters of secondary roots, and in the number of secondary roots per cluster was observed for *erf109* when compared to wildtype Col-0 (Fig. 7). Altogether, we concluded that ERF109-dependent secondary root formation upon *H. schachtii* infection relies at least partially on local auxin biosynthesis.

**Fig. 7.**
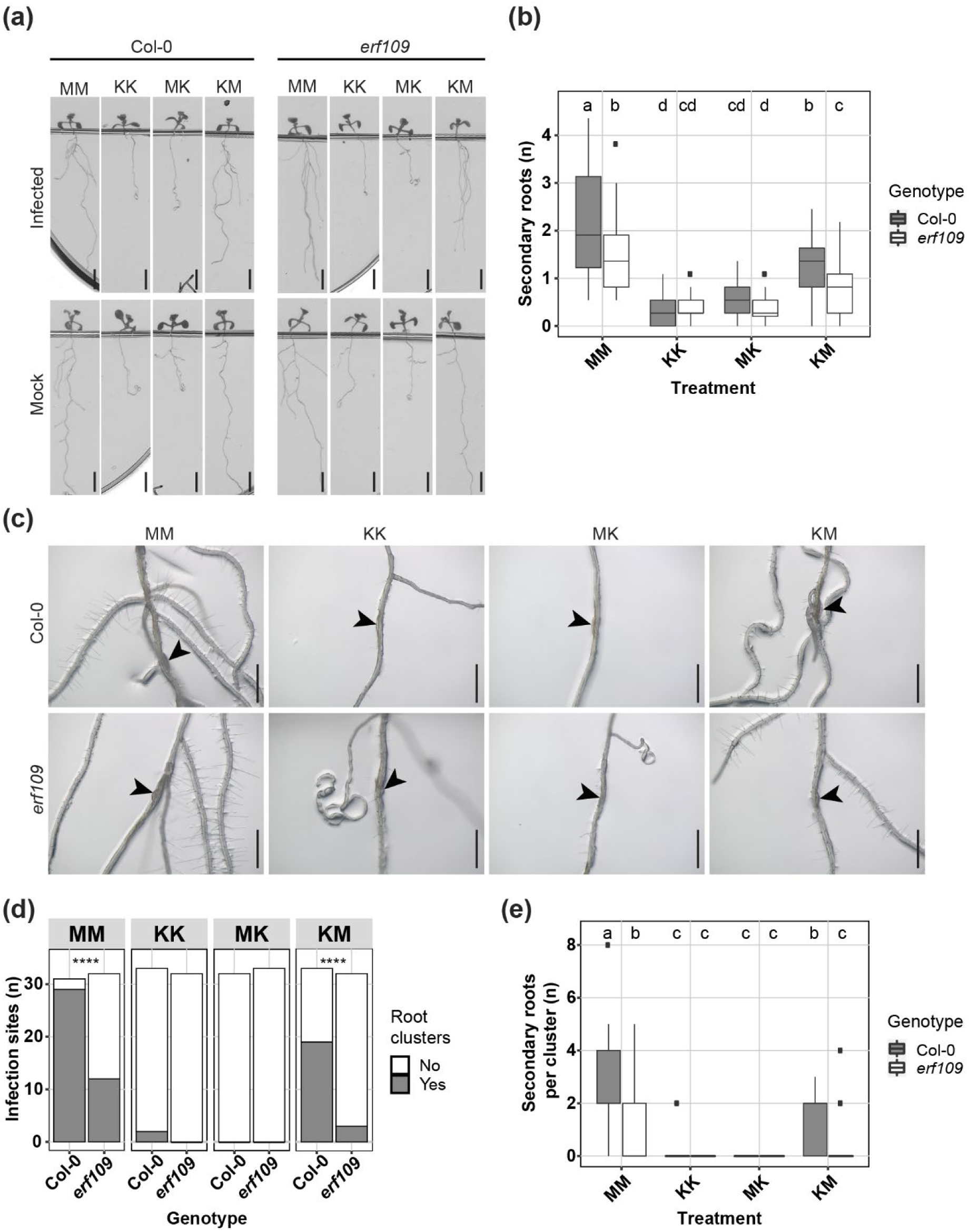
ERF109-dependent local auxin biosynthesis regulates secondary root formation upon *Heterodera schachtii* infection. Four-day-old Arabidopsis Col-0 and *erf109* seedlings were either infected with 15 *H. schachtii* J2s or mock inoculated. At 16 hours post inoculation, seedlings were transferred to treatment plates. Four treatment combinations were prepared: MM (modified Knop medium and 0.02% DMSO), KK (modified Knop medium, 10µM L-kyn and 0.02% DMSO), MK (L-kyn only in the root), KM (L-kyn only in the shoot). At 7 dpi, scans were made of the root systems and the total number of secondary roots per plant was counted. Additionally, the presence of clusters and the number of secondary roots per cluster was scored. (a) Representative pictures of wildtype Col-0 and *erf109* mutant seedlings. (b) Number of secondary roots in infected versus non-infected roots of wildtype Col-0 and *erf109* seedlings. Data of two independent biological repeats of the experiment was combined. Significance of differences in secondary roots between the different treatment combinations was calculated by ANOVA followed by Tukey’s HSD test for multiple comparisons (n=43-45, *P*<0.05). (c) Representative images of nematode infection sites in wildtype Col-0 and *erf109* mutant. (d) Number of secondary root clusters that are associated with *H. schachtii* infection sites. Data of two independent biological repeats of the experiment was combined. Statistical significance was calculated by a Pairwise Z-test n=31-33, ****, *P*<0.0001). (e) Number of secondary roots per cluster. Data of two independent biological repeats of the experiment was combined. Significance of differences between secondary roots within a cluster was calculated by Aligned Rank Transform for non-parametric factorial ANOVA followed by Tukey’s HSD test for multiple comparisons (n=31-33, *P*<0.0001). Difference in letters indicates statistically different groups. Black arrowheads indicate the infection site. Scale bar is 0.5 cm.

## Discussion

Root architecture plasticity in response to stress by soil-borne pathogens and pests is a largely unexplored field of research. Root parasitism by cyst nematodes is often associated with formation of secondary roots in proximity of infection sites (Grymaszewska & Golinowski, 1991; Goverse *et al*., 2000; Lee *et al*., 2011). However, the molecular mechanisms regulating secondary root formation in response to cyst nematode infection have thus far remained unclear. Here, we provide evidence for a model wherein formation of secondary roots near *H. schachtii* infection sites is triggered by tissue damage caused by nematode invasion. This response is regulated by the JA-dependent ERF109-activated local biosynthesis of auxin.

Our data demonstrates that secondary root formation is most likely initiated by tissue damage brought about by cyst nematode infections. The number of secondary roots induced by *H. schachtii* correlated positively with the number of nematodes that penetrated the roots. This increase in the number of secondary roots may be simply due to an increase in the number of infection sites. However, successful penetration by one J2 at a particular site often attracts other J2s, resulting in infection sites harboring multiple nematodes (Wyss & Zunke, 1986). Consistently, we also observed more nematodes within infection sites at higher inoculation densities, which correlated well with the number of secondary roots per infection site. This may mean that infection sites containing multiple nematodes developed a higher number of secondary roots per cluster compared to single-nematode associated infection sites. Moreover, we saw more extensive root tissue damage (i.e., root discoloring) at infection sites harboring multiple nematodes. We therefore consider tissue damage by infective juveniles inside roots as the likely cause of enhanced local secondary root formation.

Tissue damage in Arabidopsis leaf explants triggers *de novo* root organogenesis in a JA-dependent manner (Zhang *et al*., 2019). We found that intracellular host invasion by *H. schachtii* transiently induces JA biosynthesis and signaling, and that the JA receptor mutant *coi1-2* is defective in secondary root formation upon *H. schachtii* infection. Our results are in line with whole transcriptome analyses of root segments of Arabidopsis harboring migrating juveniles of *H. schachtii* at 10 hpi, which also showed that JA biosynthesis and signaling genes are upregulated during host invasion (Kammerhofer *et al*., 2015; Mendy *et al*., 2017). In contrast, recent reports indicate that host invasion by *H. schachtii* does not activate the JA signaling biosensor *JAZ10::NLS-3xVENUS* in Arabidopsis roots (Marhavy *et al*., 2019). The discrepancy between our observations with the *JAS9-VENUS* biosensor and the observations with the *JAZ10::NLS-3xVENUS* biosensor might be due to differences in sensitivity of both sensor constructs. As compared to *JAZ10::NLS-3xVENUS,* the *JAS9-VENUS* biosensor is particularly sensitive to biologically active JA (JA-isoleucine) enabling the visualization of local JA signaling in response to stress in Arabidopsis roots at a high spatiotemporal resolution (Larrieu *et al*., 2015). Furthermore, *JAS9-VENUS* has been used to monitor the dynamics of JA signaling in response to single cell ablation and intercellular migration of the less-damaging root-knot nematodes in Arabidopsis roots (Zhou *et al*., 2019). Therefore, based on the activity of the *JAS9-VENUS* biosensor in our experiments, we conclude that the tissue damage associated with host invasion triggers a JA signal in cells close to the infection site of *H. schachtii*. Moreover, the transient nature of the JA signal suggests that the damage trigger decreases after nematode host invasion, or that JA signaling is actively suppressed by *H. schachtii* when infective juveniles become sedentary.

JA signaling during *H. schachtii* migration also results in activation of plant defense responses (Kammerhofer *et al*., 2015). We observed that the *coi1-2* mutant is more susceptible to penetration by *H. schachtii*, which is in line with previous findings showing a negative effect of exogenous JA on *H. schachtii* penetration rate (Kammerhofer *et al*., 2015). However, after nematode penetration, COI1 does not affect the rate at which J2s induce a permanent feeding site (Marhavy *et al*., 2019). Altogether, these findings suggest that JA signaling both negatively regulates host penetration rate by *H. schachtii* and mediates secondary root formation at *H. schachtii* infection sites.

The damage-induced formation of secondary roots by *H. schachtii* appears to be regulated by the JA-dependent expression of *ERF109*. We found that the expression of *ERF109*, which showed the same transient induction pattern as the JA biosynthesis reporter *AOS* and *JAS9-VENUS* biosensor, was abrogated in *coi1-2* mutant. Moreover, the *erf109* mutant was as defective as the *coi1-2* mutant in the density-dependent secondary root formation upon *H. schachtii* infection. Consistently with our data, *ERF109* expression showed a COI1-dependent transient expression upon wounding in leaf explants (Zhang *et al*., 2019). Furthermore, the *erf109* mutation also disrupted the induction of secondary root formation by exogenous application of JA (Cai *et al*., 2014). Altogether, our findings show that tissue damage by invading nematodes triggers a JA signal, which induces the ERF109-dependent formation of secondary roots.

Next, our data provides evidence that damage-induced activation of *ERF109* regulates formation of secondary roots via local auxin biosynthesis. The local accumulation of auxin at nematode infection sites (i.e., expression of the auxin reporter *DR5::GUS*) was strongly reduced in the *erf109* mutant compared to wildtype plants. However, when auxin biosynthesis was blocked in whole seedlings or only in roots, the local accumulation of auxin at nematode infection sites was completely abolished in both the *erf109* mutant and wildtype Arabidopsis. Taken together, this demonstrates that auxin accumulation at nematode infection sites is at least partially dependent on ERF109-regulated local auxin biosynthesis. Importantly, the patterns observed for local accumulation of auxin at nematode infection sites matched the patterns of secondary root formation in absence or presence of the auxin biosynthesis inhibitor. The inhibition of auxin biosynthesis in the roots, but not in the shoots, abolished the formation of secondary roots upon nematode infection. Previously, ERF109 was shown to regulate secondary root formation by binding the promoter of auxin biosynthesis genes upon exogenous application of JA (Cai *et al*., 2014). Here, our data shows that tissue damage by nematodes activates JA signaling and subsequently induces ERF109, which on its turn regulates secondary root formation via local biosynthesis of auxin.

After blocking auxin biosynthesis in the shoots, we observed auxin accumulation and formation of secondary roots at nematode infection sites, which indicates that polar auxin transport from the shoots is not required for secondary root formation at nematode infection sites. Nevertheless, we noted a quantitative effect of the inhibition of auxin biosynthesis in shoots, leading to the formation of fewer secondary root clusters and less secondary roots per cluster as compared to untreated plants. This implicates that polar auxin transport from the shoots may still play a complementary role in secondary root formation at nematode infection sites, albeit below the detection levels of the *DR5::GUS* reporter. Polar auxin transport from the shoots and further redistribution in root tissue results from the coordinated activities of auxin influx and efflux carrier proteins (Petrasek & Friml, 2009). Lee *et al*. (2011) showed that *H. schachtii* induces the formation of secondary roots in double *aux1lax3* and quadruple *aux1lax1lax2lax3* influx carrier mutants, which are otherwise unable to form secondary roots. This suggests that the accumulation of auxin and subsequent formation of secondary roots may be regulated independently of the activity of these influx carriers. There is ample evidence that auxin efflux carriers (i.e., PIN proteins) are important for the susceptibility of Arabidopsis to infections of *H. schachtii* (Grunewald *et al*., 2009). However, if and how they might contribute to the accumulation of auxin underlying the damage-induced formation of secondary roots needs further investigation.

Here, we demonstrate that ERF109-mediated local adaptations in root architecture compensate for primary root growth inhibition in response to nematode infection. In wildtype Arabidopsis, increasing densities of J2s led to a decline in the length of infected primary roots. However, this reduction in length of infected primary roots did not result into a smaller root system, because of an increase in the total length of secondary roots. Our data shows that these adaptations in root architecture depend on the transient and local activation of *ERF109* by JA at nematode infection sites. Consistently, the JA signaling mutant *coi1-2* showed a similar impairment as *erf109* in compensating primary root length inhibition by an increase in total secondary root length (Fig. S5). Nevertheless, since COI1 also affects plant susceptibility to nematode penetration more complex defense versus growth trade-offs may influence root growth in the *coi1-2* mutant. Importantly, loss of function mutations in *ERF109* do not alter the susceptibility of Arabidopsis to *H. schachtii* penetration, but instead affect root architecture plasticity in response to nematode infection. Further research is needed to understand whether ERF109-mediated compensatory adaptations in root architecture could mediate tolerance of Arabidopsis to infections by *H. schachtii*.

It was previously shown that meristem damage caused by *M. incognita* root tip penetration triggers regeneration via JA- and ERF109-mediated damage signaling (Zhou *et al*., 2019). Here, we show that *H. schachtii* penetration of the mature root zone causes damage-induced secondary root formation, which compensates for primary root growth inhibition. Therefore, we consider root tip regeneration and secondary root formation as two different outcomes of the same compensatory mechanism in response to tissue damage in different root zones.

Furthermore, we show the first case of a naturally occurring and biotic stress that triggers damage signaling-mediated secondary root formation. Primary roots can form two types of secondary roots (Sheng *et al*., 2017). One type, referred to as lateral root, forms during the physiological post-embryogenic development of plants and is regulated by ARF7 and ARF19 auxin response factors. The other type is induced by sterile mechanical injury of the mature root zone, soil penetration, or osmotic stress, and is dependent on the transcription factor WOX11. Sterile mechanical injury causes a different type of root tissue damage compared to a biotic stress such as cyst nematodes (Marhavy *et al*., 2019). Sterile mechanical injury damages many root cells at one time. Instead, cyst nematode host invasion causes the rupture of multiple single cells one after the other over the course of many hours (Wyss & Zunke, 1986). Thus, our results provide biological relevance for a mechanism so far only observed upon artificial conditions.

As a natural trigger for damage signaling, *H. schachtii* can be used to further elucidate the pathway leading to secondary root formation. ERF109 was previously found to be responsive to reactive oxygen species (ROS) (Kong *et al*., 2018). It would be interesting to test whether ROS mediate ERF109-dependent secondary root formation upon *H. schachtii* infection. Furthermore, follow up research could investigate if damage receptors activated during *H. schachtii* migration (Shah *et al.,* 2017) act upstream of ERF109. The root-knot nematode *M. javanica* triggers expression of LBD16, a downstream target of both WOX11, and ARF7 and ARF19 (Cabrera *et al*., 2014; Olmo *et al*., 2017). Moreover, *M. javanica* infection of primary roots induces secondary root formation independently from ARF7 and ARF19 (Olmo *et al*., 2017). This suggests that nematode-induced secondary root formation could be regulated by WOX11. However, whether WOX11-mediated secondary root formation acts downstream of the ERF109-damage signaling pathway remains unknown.

In summary, we showed that *H. schachtii* triggers the formation of secondary roots via JA- and ERF109-mediated damage signaling (Fig. 8). Furthermore, ERF109-mediated secondary root formation compensates for primary root growth inhibition associated with *H. schachtii* infection. Thus, damage signaling-induced formation of secondary roots points at a novel mechanism underlying plant root architecture plasticity to biotic stress. Further research is needed to investigate whether damage-induced root architecture plasticity can contribute to plant tolerance to belowground herbivory.

**Fig. 8.**
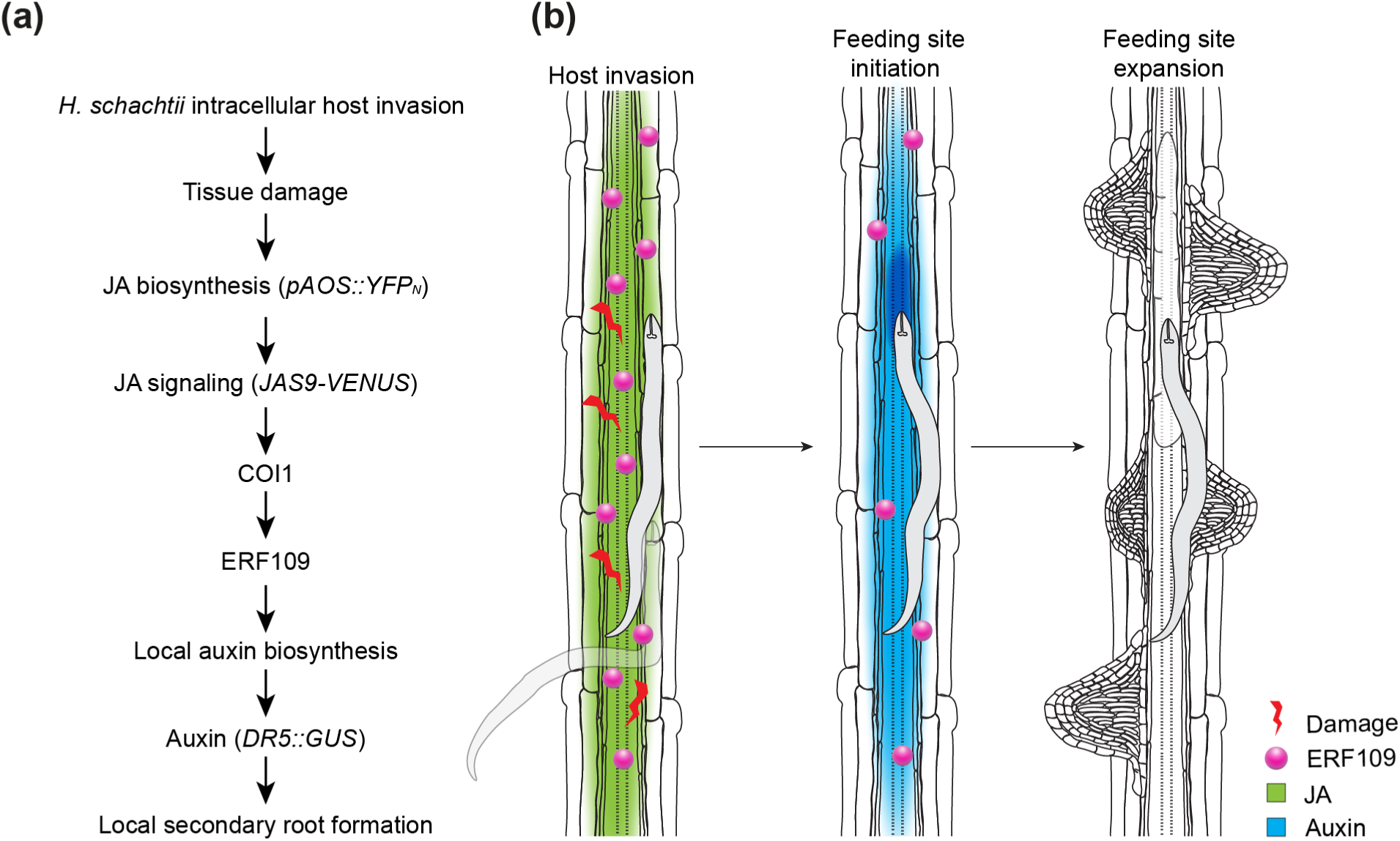
Model of the pathway regulating *Heterodera schachtii*-induced secondary root formation. (a) Intracellular invasion of host roots by *H. schachtii* causes tissue damage, which triggers JA biosynthesis (*pAOS::YFPN*). JA signaling (*JAS9-VENUS*) via COI1 induces expression of *ERF109*, which leads to auxin accumulation (*DR5::GUS*) via local auxin biosynthesis. ERF109-mediated local auxin biosynthesis finally results in the formation of secondary roots at *H. schachtii* infection sites. (b) Graphical model illustrating the pathway investigated in this paper.

## Supporting information

Supporting information S1toS6

## Acknowledgements

We thank Hang Liu for the help provided with data collection as part of his MSc Thesis at Wageningen University. This work was supported by the Graduate School Experimental Plant Sciences (EPS). WZ is funded by EMBO long-term fellowship (ALTF 784-2014) and the National Natural Science Foundation of China (32070874). JJW is funded by Dutch Top Sector Horticulture & Starting Materials (TU18152). MGS was supported by NWO domain Applied and Engineering Sciences VENI grant (17282). JLLT was supported by NWO domain Applied and Engineering Sciences VENI (14250) and VIDI (18389) grants. No conflict of interest declared.

## Author contributions

JLLT, GS, NG, JJW, AG, WZ, and VW conceived the project. NG, JJW and MSH designed the experiments and performed data collection. WZ and VW provided most of the Arabidopsis mutants and granted access to the confocal microscope. WZ performed the crossing to obtain the *pERF109::GFP/coi1-2* Arabidopsis line, while homozygous plants were selected by both WZ and NG. Data was analyzed and interpreted by NG, JJW, MSH and MGS. NG, JLLT, GS, and JJW wrote the article with inputs from AG, MGS, VW, WZ, MSH and FMWG.

## Data availability

The data that support the findings of this study is available from the corresponding author upon request.

## Supporting information

Supporting Information Fig. S1. Quantification of *pERF109::GFP* nuclear fluorescence intensity in infected wildtype Col-0 and *coi1-2* mutant.

Supporting Information Fig. S2. *DR5::GUS* saturation in root tips of infected wildtype Col-0 and *erf109* mutant seedlings under four different treatment combinations with and without L-kyn applied to shoots and/or roots.

Supporting Information Fig. S3. Number of secondary roots in non-infected roots of seedlings of wildtype Col-0 and *erf109* mutant under four different treatment combinations with and without L-kyn applied to shoots and/or roots.

Supporting Information Fig. S4. Root architecture of uninfected *coi1-2* and *erf109* Arabidopsis plants compared to wildtype Col-0 plants.

Supporting Information Fig. S5. COI1-mediated secondary root formation allows for maintenance of total root length despite primary root growth inhibition by *Heterodera schachtii*.

Supporting Information Fig. S6. Yuc split plate assay showing that ERF109 regulates local auxin biosynthesis at the nematode infection site.

## Notes

### Competing Interest Statement

The authors have declared no competing interest.

