## Supporting information S1toS6 for "Root architecture plasticity in response to endoparasitic cyst nematodes is mediated by damage signaling"

The following Supporting Information is available for this article:

**Fig. S1** Quantification of *pERF109::GFP* nuclear fluorescence intensity in infected wildtype Col-0 and *coi1-2* mutant.

**Fig. S2** *DR5::GUS* saturation in root tips of infected wildtype Col-0 and *erf109* mutant seedlings under four different treatment combinations with and without L-kyn applied to shoots and/or roots.

**Fig. S3** Number of secondary roots in non-infected roots of seedlings of wildtype Col-0 and *erf109* mutant under four different treatment combinations with and without L-kyn applied to shoots and/or roots.

**Fig. S4** Root architecture of uninfected *coi1-2* and *erf109* Arabidopsis plants compared to wildtype Col-0 plants.

**Fig. S5** COI1-mediated secondary root formation allows for maintenance of total root length despite primary root growth inhibition by *Heterodera schachtii*.

**Fig. S6** Yuc split plate assay showing that ERF109 regulates local auxin biosynthesis at the nematode infection site.

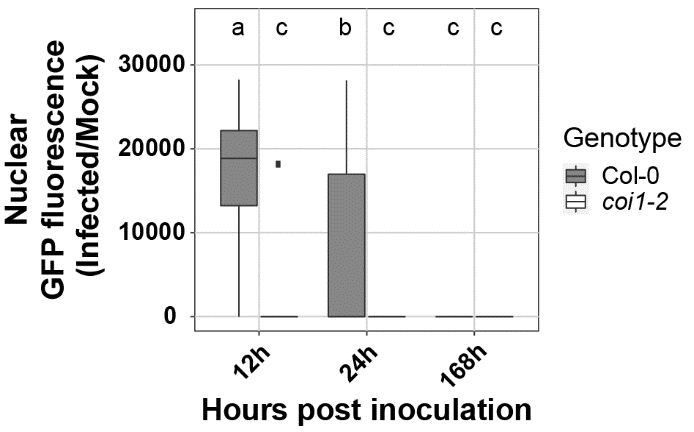

**Fig. S1** Quantification of *pERF109::GFP* nuclear fluorescence intensity in infected wildtype Col-0 and *coi1-2* mutant. Four-day-old Arabidopsis seedlings were either inoculated with 15 *H. schachtii* J2s or mock inoculated. At 12, 24, and 168 hpi seedlings were mounted in 10 μg ml^-1^ propidium iodide (PI) and then imaged. Nuclei were selected and the integrated density was measured using Fiji software. Ratio of the nuclear GFP fluorescence between infected and non-infected seedlings of wildtype Col-0 and *coi1-2* mutant. Significance of differences between fluorescent intensity in Col-0 and *coi1-2* roots over the different timepoints was calculated by Aligned Rank Transform for non-parametric factorial ANOVA followed by Tukey’s HSD test for multiple comparisons (n=30, *P*<0.001). Difference in letters indicates statistically different groups.

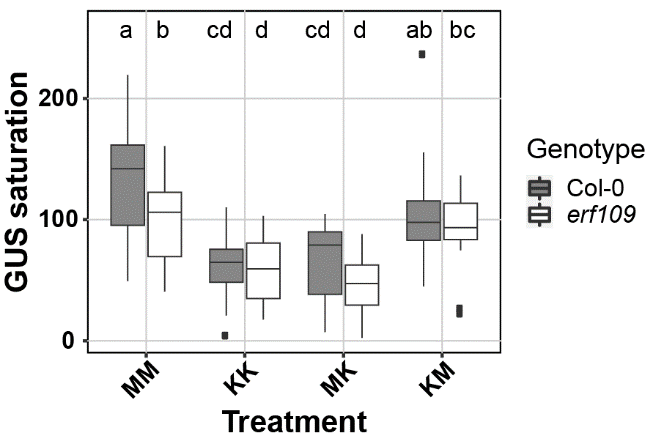

**Fig. S2** *DR5::GUS* saturation in root tips of infected wildtype Col-0 and *erf109* mutant seedlings under four different treatment combinations with and without L-kyn applied to shoots and/or roots. Four-day-old wildtype Arabidopsis Col-0 and *erf109* mutant seedlings expressing the auxin *DR5::GUS* reporter were infected with 15 *H. schachtii* J2s. At 16 hours post inoculation, seedlings were transferred to treatment plates in a split-plate design. Four treatment combinations were prepared in split-plate assay: MM (modified Knop medium and 0.02%DMSO), KK (modified Knop medium, 10µM L-kyn and 0.02% DMSO), MK (L-kyn only in the root), KM (L-kyn only in the shoot). At 3 dpi GUS staining was performed for 4 hours and seedlings were imaged. GUS saturation was measured as mean grey value using Fiji software. Data of two independent biological replicates was combined. Significance was calculated by ANOVA followed by Tukey’s HSD test (n=20, *P*<0.05). Difference in letters indicates statistically different groups.

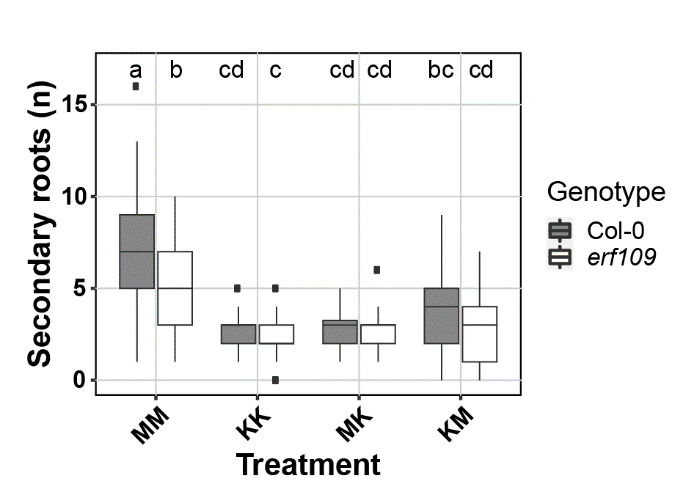

**Fig. S3** Number of secondary roots in non-infected roots of seedlings of wildtype Col-0 and *erf109* mutant under four different treatment combinations with and without L-kyn applied to shoots and/or roots. Four-day-old Arabidopsis wildtype Col-0 and *erf109* seedlings were either infected with 15 *H. schachtii* J2s or mock inoculated. At 16 hours post inoculation, seedlings were transferred to treatment plates in a split-plate design. Four treatment combinations were prepared: MM (modified Knop medium and 0.02% DMSO), KK (modified Knop medium, 10µM L-kyn and 0.02% DMSO), MK (L-kyn only in the root), KM (L-kyn only in the shoot). At 7 dpi, scans were made of the root systems and the total number of secondary roots per plant was counted. Data of two independent biological repeats of the experiment was combined. Significance of differences in secondary roots between the different treatment combinations was calculated by ANOVA followed by Tukey’s HSD test for multiple comparisons (n=43-45, *P*<0.001). Difference in letters indicates statistically different groups.

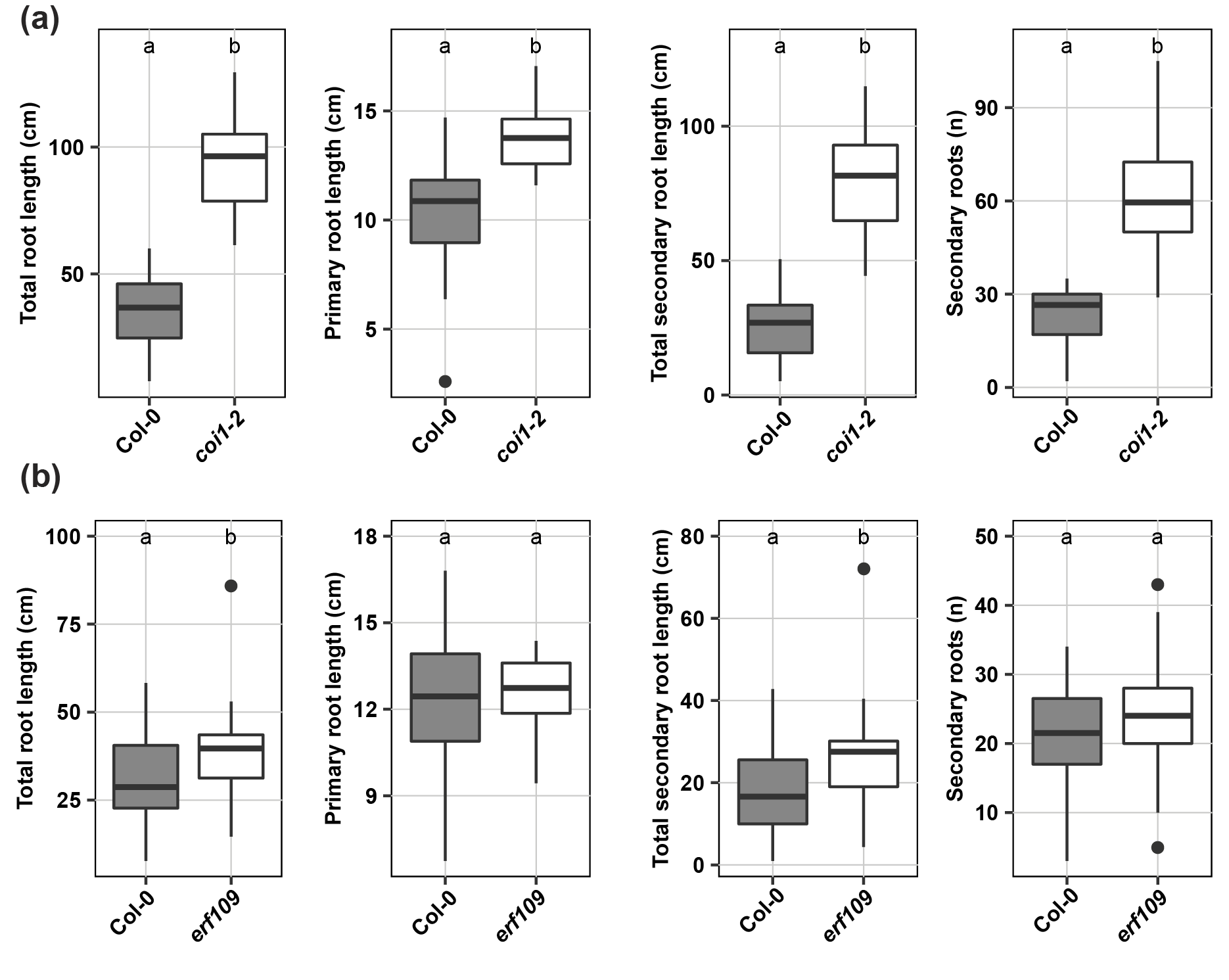

**Fig. S4** Root architecture of uninfected *coi1-2* and *erf109* Arabidopsis plants compared to wildtype Col-0 plants. Scans of the roots of 16-day-old plants were made and the total root length was measured using WinRHIZO. For the experiment including *coi1-2* (a) the primary root was measured manually using ImageJ because of the complex root system of the mutant. For the experiment including *erf109* (b) the primary root length was automatically measured by WinRHIZO. Total secondary root length was calculated by subtracting the primary root length from the total root length. Data from three independent biological repeats of the experiment was combined. Significance of differences between genotypes was calculated by Student’s T test (n=30, *P*<0.05). ). Different letters indicate statistically different groups.

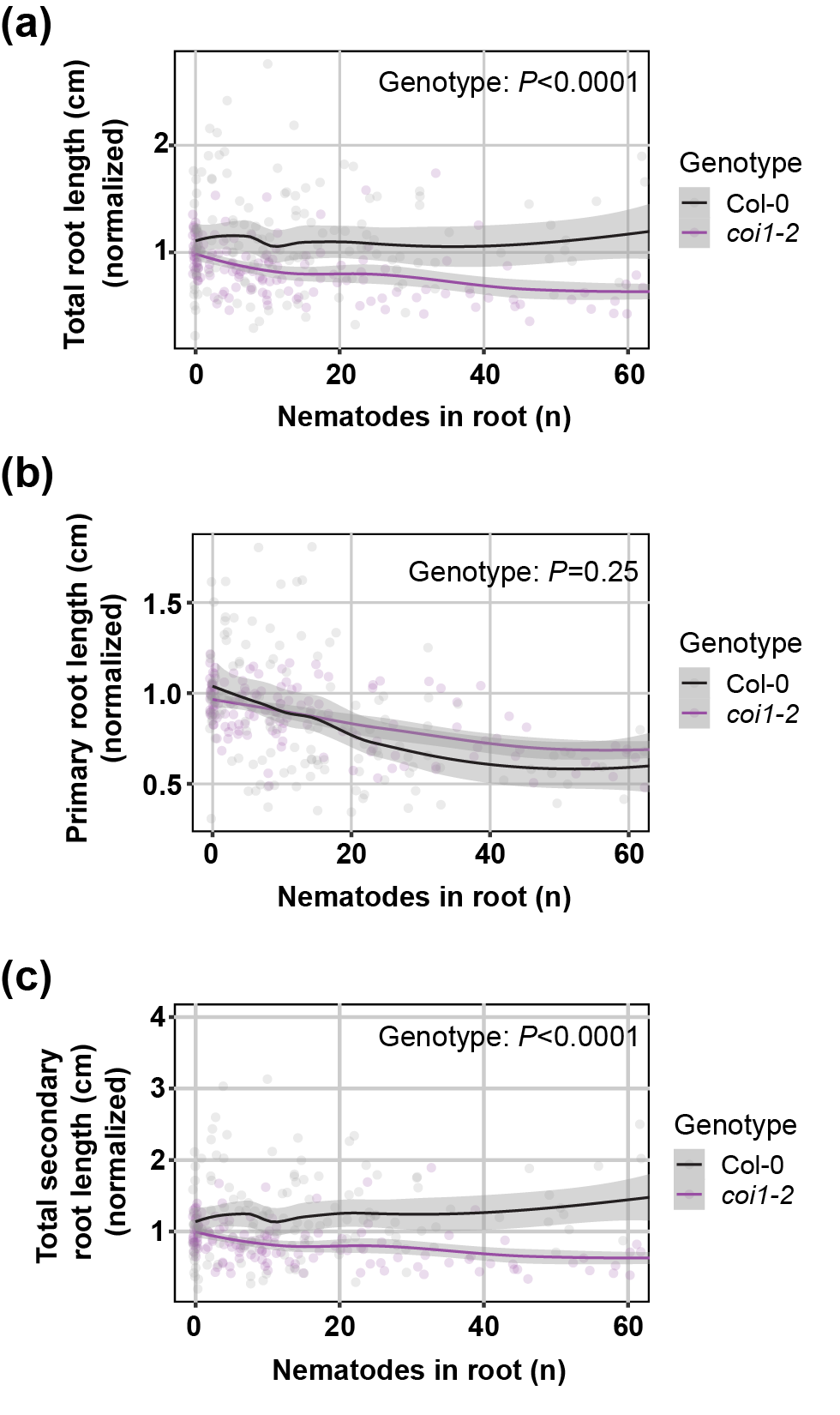

**Fig. S5** COI1-mediated secondary root formation allows for maintenance of total root length despite primary root growth inhibition by *Heterodera schachtii*. Nine-day-old Col-0 and *coi1-2* Arabidopsis seedlings were inoculated with increasing *H. schachtii* densities ranging from 0 (mock) to 500 J2s per seedling. At 7 dpi, scans were made of the root systems and the root length was measured using WinRHIZO. Total, primary, and secondary root length was normalized to the average respective component in mock-treated roots. Fuchsin staining was performed for counting the number of J2s that penetrated the roots. (a) Total root length per number of nematodes in the roots. (b) Primary root length per number of nematodes in the roots. (c) Total secondary root length per number of nematodes in the roots. Data from three independent biological repeats of the experiment was combined. Significance of differences between genotypes was calculated by ANOVA (n=30). Grey area indicates the 95% confidence interval.

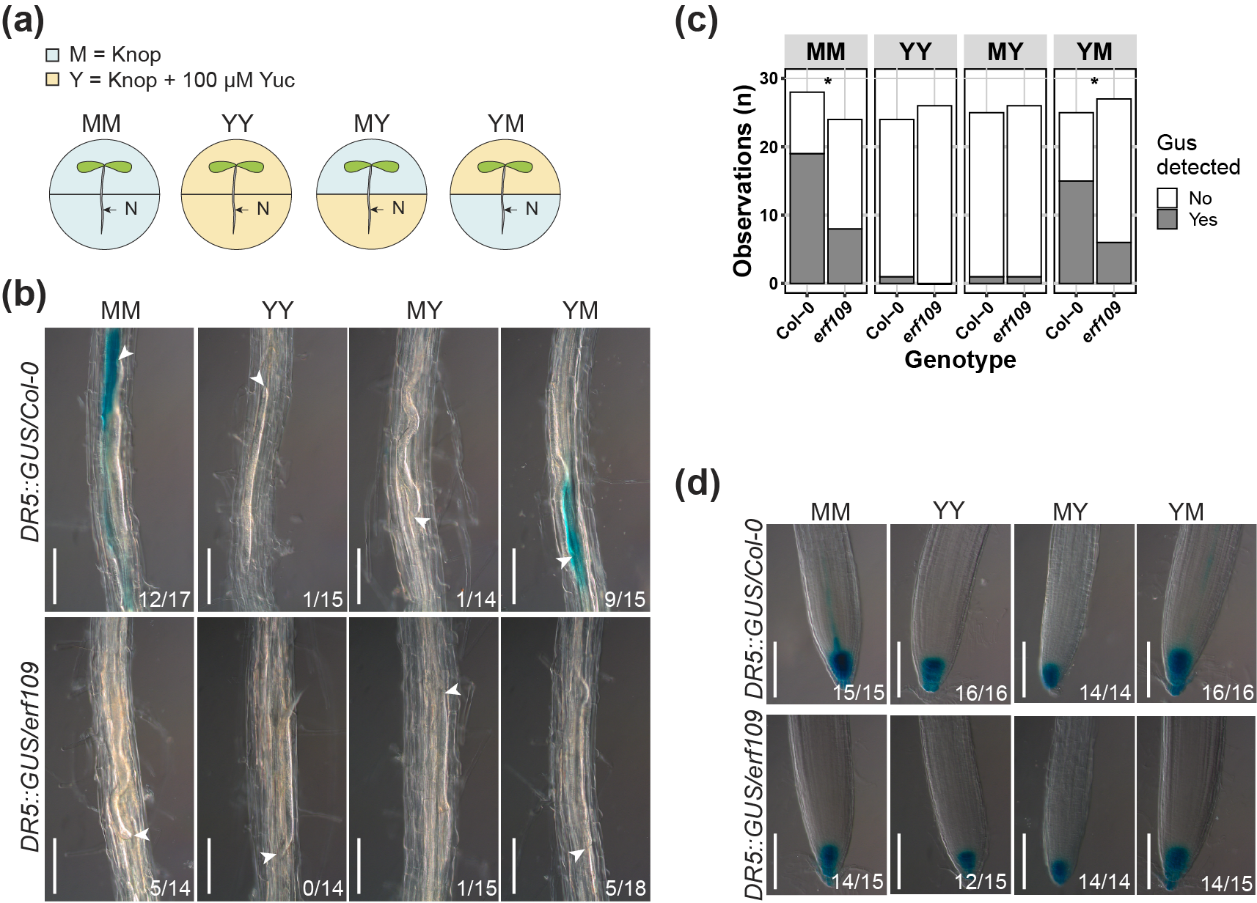

**Fig. S6** Yuc split plate assay showing that ERF109 regulates local auxin biosynthesis at the nematode infection site. Four-day-old Arabidopsis Col-0 and *erf109* seedlings expressing the auxin *DR5::GUS* reporter were infected with 15 *Heterodera schachtii* J2s. At 16 hours post inoculation, seedlings were transferred to treatment plates. Four treatment combinations were prepared: MM (modified Knop medium and 0.2% DMSO), YY (modified Knop medium, 100µM Yuc and 0.2% DMSO), MK (Yuc only in the root), YM (Yuc only in the shoot). At 3 dpi GUS staining assay was performed for 4 hours and seedlings were imaged. Single-nematode infection sites were selected for observation. (a) Experimental design with Arabidopsis seedlings transferred to split plates with modified Knop medium either with or without Yuc. N = nematode. (b) *DR5::GUS* expression at nematode infection sites in wildtype Col-0 and *erf109* roots in the four different treatment combinations with or without Yuc applied to shoots and/or roots. (c) Number of observations with (Yes) or without (No) GUS staining at the nematode infection sites in roots of wildtype Col-0 and *erf109* plants. Statistical significance was calculated by a Pairwise Z-test (n=33, *, *P*<0.05). (d) *DR5::GUS* expression in the root tips of Col-0 and *erf109* roots. White arrowheads indicate the nematode head. Frequencies at the bottom right corner indicate how many times GUS staining was observed in one of the three independent biological repeats of the experiment. Scale bar is 200 µm.
